# Density dependence during evolutionary rescue increases extinction risk but does not prevent adaptation

**DOI:** 10.1101/2024.12.12.628101

**Authors:** Laure Olazcuaga, Brett. A Melbourne, Scott W. Nordstrom, Ruth A Hufbauer

## Abstract

Evolutionary rescue allows populations to adapt and persist despite severe environmental change. While well studied under density-independent conditions, the role of density dependence, including competition, remains unclear. Theoretical models offer conflicting predictions, with density dependence either increasing extinction risk or enhancing adaptation. We empirically tested how density dependence influences evolutionary rescue by exposing experimental populations to a stressful environment for six generations under density-dependent or independent conditions, with populations where either evolution was possible or was prevented by replacing individuals each generation. Density dependence suppressed population size and increased extinction risk, whereas density independence enabled rapid growth, especially in genetically diverse populations where evolution was possible. Although density dependence raises extinction risk, it does not prevent populations from responding to selection, since surviving density-dependent populations still exhibited increased intrinsic and realized fitness. These findings reconcile theoretical discrepancies, showing density dependence can simultaneously increase extinction risk but may favor adaptation. Our results underscore the importance of considering density dependence in conservation strategies.

## Introduction

Human-altered environments are leading to drastic declines in biodiversity globally (Ceballos *et al*. 2017; Diaz *et al*. 2018). Conservation interventions can be effective (Langhammer *et al*. 2024), but are unlikely to be sufficient on their own (Senior *et al*. 2024). In addition to the need for fundamental changes to human resource use (Diaz *et al*. 2018), it is vital that we also understand the conditions under which species will be able to adapt to human-altered environments.

Evolutionary rescue is adaptation that enables populations to persist and grow following habitat alteration severe enough that it otherwise would cause extinction (Gomulkiewicz & Holt 1995). Prior to environmental stress, population sizes in nature are limited by multiple abiotic and biotic factors. When severe environmental stress occurs that reduces population growth rates below the replacement rate, population size will decline, potentially to extinction. If adaptation to new conditions is rapid enough, populations will be able to grow again. Evolutionary rescue thus entails a U-shaped curve in population size through time, with an adaptive increase in fitness that prevents extinction.

Theory and multiple experiments have demonstrated that evolutionary rescue can enable population persistence, with the probability of rescue increasing with genetic diversity (Bell 2013; Bell & Gonzalez 2009; Carlson *et al*. 2014; Hufbauer *et al*. 2015; Olazcuaga *et al*. 2023; Orr & Unckless 2008; Ramsayer *et al*. 2013).

However, in most of that research, density-independent growth is assumed theoretically, or populations are maintained at low density with low intraspecific competition for resources or other forms of negative density dependence (e.g., Hufbauer et al. 2015, Lewis et al. 2024). However, density dependence is nearly ubiquitous in nature and thus it is crucial to study how density dependence influences the process of evolutionary rescue. Additionally, environmental change can reduce carrying capacity, intensify competition or alter trait–fitness landscapes, thereby modifying ecological constraints and selective pressures, making it even more important to evaluate the role of density dependence in the context of evolutionary rescue.

While positive density dependence can occur in nature, e.g., through Allee effects, which lower fitness at small population size due to difficulty finding mates or engaging in social cooperative behavior (Kramer et al. 2009, Muir et al. 2024), we are primarily concerned with negative density dependence, where population performance drops as abundance increases. Negative density dependence is frequent when habitat size is restricted, which is common for populations of conservation concern in the modern era of environmental degradation and habitat fragmentation (Scholl *et al*. 2022). Given this, the findings that evolutionary rescue occurs when habitats are relatively large or resources relatively abundant (e.g., Agashe *et al*. 2011; Bell & Gonzalez 2009; Hufbauer *et al*. 2015) cannot be confidently applied to many situations. Thus, how density dependence affects evolutionary rescue represents a key knowledge gap in both fundamental understanding of the rescue process, and how to enhance evolutionary rescue in nature.

Theoretical models have addressed this knowledge gap by comparing outcomes between populations experiencing density independence and those experiencing density dependence, typically negative density dependence (Chevin & Lande 2010; Nordstrom *et al*. 2023; Orr & Unckless 2008; Reed *et al*. 2015; Uecker *et al*. 2014; Vinton & Vasseur 2020). However, their findings are contradictory. Some studies show that density dependence hinders rescue through its effects on demography, as it renders populations more vulnerable to extinction due in part to a faster initial decline and slower recovery in population size (Chevin & Lande 2010; Nordstrom *et al*. 2023; Orr & Unckless 2008; Uecker *et al*. 2014). However, others find that density dependence can reduce extinction risk through ecological release that leads to an increase in fecundity or decrease in mortality when population size drops (Reed *et al*. 2015; Vinton & Vasseur 2020). These contradicting predictions stem from differences in model assumptions, including whether models start with equivalent intrinsic fitness (e.g., Chevin & Lande 2010; Nordstrom *et al*. 2023; Orr & Unckless 2008; Uecker *et al*. 2014) or equivalent realized fitness (Reed *et al*. 2015; Vinton & Vasseur 2020) of populations under density dependence and density independence. While intrinsic fitness refers to the potential contribution of individuals to the population growth rate in the absence of density effects, realized fitness refers to the observed growth rate, which is additionally influenced by density (i.e. an individual’s fitness is modified by intraspecific interaction). Models that assume equivalent realized fitness prior to environmental change create a situation where populations adapted to high density will have increased growth when negative density dependence is relaxed following environmental change. This distinction is critical, as starting from equal intrinsic versus equal realized fitness can lead to contrasting expectations for the likelihood of rescue. Data from empirical systems will be useful to shed light on how density dependence alters extinction risk in nature.

Most theoretical studies of the role of density dependence focus on how the demographic effects of density dependence influence the probability of rescue (or its inverse, extinction) shortly after environmental change, but not on the evolutionary effects on fitness of the surviving populations after rescue. However, fitness following evolutionary rescue is crucial to understanding how populations might respond in the long term to environmental change. Persisting populations adapting under density dependence may have higher fitness (compared to persisting populations adapting under density independence) through at least two mechanisms. First, theory shows that if density dependence and the change in the environment select for similar phenotypes, the speed and/or magnitude of adaptation and the eventual fitness achieved in the new environment will be higher than under density independence (Osmond & de Mazancourt 2013). Second, because density dependence can lead to higher extinction rates, the populations that persist can on average have higher fitness after rescue due to survivorship bias, as those with lower fitness are more likely to go extinct (Nordstrom *et al*. 2023). Thus, even if density dependence reduces the probability of rescue by increasing extinction risk, it may also benefit surviving populations through increased fitness. These effects of density dependence on extinction and fitness may be modulated by genetic variation. For example, the demographic costs of density dependence may increase the importance of harboring sufficient genetic diversity to adapt rapidly to environmental change, and those with low diversity or high genetic load, for example because of prior bottlenecks in population size, might not be able to adapt rapidly enough.

Here, we experimentally test for the first time how density-dependent population dynamics affect evolutionary rescue following sudden environmental change. We use *Tribolium castaneum* as a model system (Pointer *et al*. 2021). In our experiment, we manipulated three factors with two treatments each: density dependence (density independent and density dependent), diversity (diverse and bottlenecked), and evolution (evolution possible or not). We tracked the dynamics of 188 experimental populations in an environment made harsh by a pesticide (Durkee *et al*. 2024) for 6 generations. In ninety-two of these populations, evolution was possible, and in the other 92 populations evolution was not possible (i.e., control populations). Through comparison with no-evolution control populations, we tracked how changes in fitness consistent with adaptation influenced ecological dynamics throughout the experiment. Additionally, at the end of the experiment, we measured population realized fitness in the stressful environment at three densities to further evaluate how surviving populations responded to the environmental stress and to population density.

This experimental set-up allowed us to address the following question: How do density dependence, bottlenecks, and evolution influence evolutionary rescue? We tracked three key hallmarks of rescue: extinction, population dynamics, and the evolution of fitness in a stressful environment. We hypothesized that when evolution was possible (1) density dependence would increase the probability of extinction due to stronger demographic constraints relative to density independence, but that (2) if such populations survived under conditions of density dependence they could develop higher fitness than density independent populations, consistent with stronger adaptation following rescue and that (3) these responses should be mediated by the genetic diversity of the populations, with more diverse populations adapting more effectively, and bottlenecked populations experiencing higher extinction risk. Further, we hypothesized that (4) populations prevented from evolving, would have higher extinction rates and no increases in fitness compared to populations where evolution was possible.

## Methods

### Model system and lab conditions

Our experiment used *Tribolium castaneum*, the red flour beetle. During laboratory maintenance and in the experiments, *T. castaneum* populations were kept in 4 cm by 4 cm by 6 cm acrylic containers with 20 g of medium (hereafter a ‘patch’), which served as both a habitat and a food source. The life cycle of *T. castaneum* was constrained to mimic that of a seasonally breeding organism with non-overlapping generations (Melbourne & Hastings 2008). Specifically, each generation was initiated by giving adult beetles fresh medium for 24 hours to reproduce. Adults then were removed from the patches, and the eggs were left to mature into adults for an additional 34 days per generation. Beetles were kept in incubators at 31°C with 54 ± 16% relative humidity. Several incubators were used, and patches were randomized among and within incubators weekly to prevent systematic incubator effects. This process of randomization avoids in particular problems linked to the spatial heterogeneity of humidity across growth chambers.

### Creation of diverse and bottlenecked populations

We started with a large, diverse stock population (see Durkee *et al*. 2023, 2024 for details). From this we created 47 independent source populations: 24 bottlenecked and 23 diverse. Each diverse source population was initiated with an independent group of 500 individuals from the stock population. Each bottlenecked source population was initiated from an independent sibling pair from the stock population, (the “strong bottleneck populations” in Olazcuaga *et al*. 2023). These 47 source populations were maintained at a size of 400-500 individuals for 8-10 generations on standard medium (95% wheat flour and 5% brewer’s yeast), distributed over four temporal blocks, one week apart.

### Evolutionary rescue experiment

We performed a multifactorial evolutionary rescue experiment manipulating density dependence (density independent vs density dependent), genetic diversity (diverse vs bottleneck), and evolution (evolution possible vs no evolution), giving eight treatment combinations (2×2×2, Fig. 1). Each of the 47 source populations (diverse and bottlenecked) was used to found 4 experimental populations with 100 individuals, giving 188 experimental populations ([23 diverse + 24 bottlenecked source populations] × 2 density treatments × 2 evolution treatments). These experimental populations were distributed in four temporal blocks.

**Figure 1.**
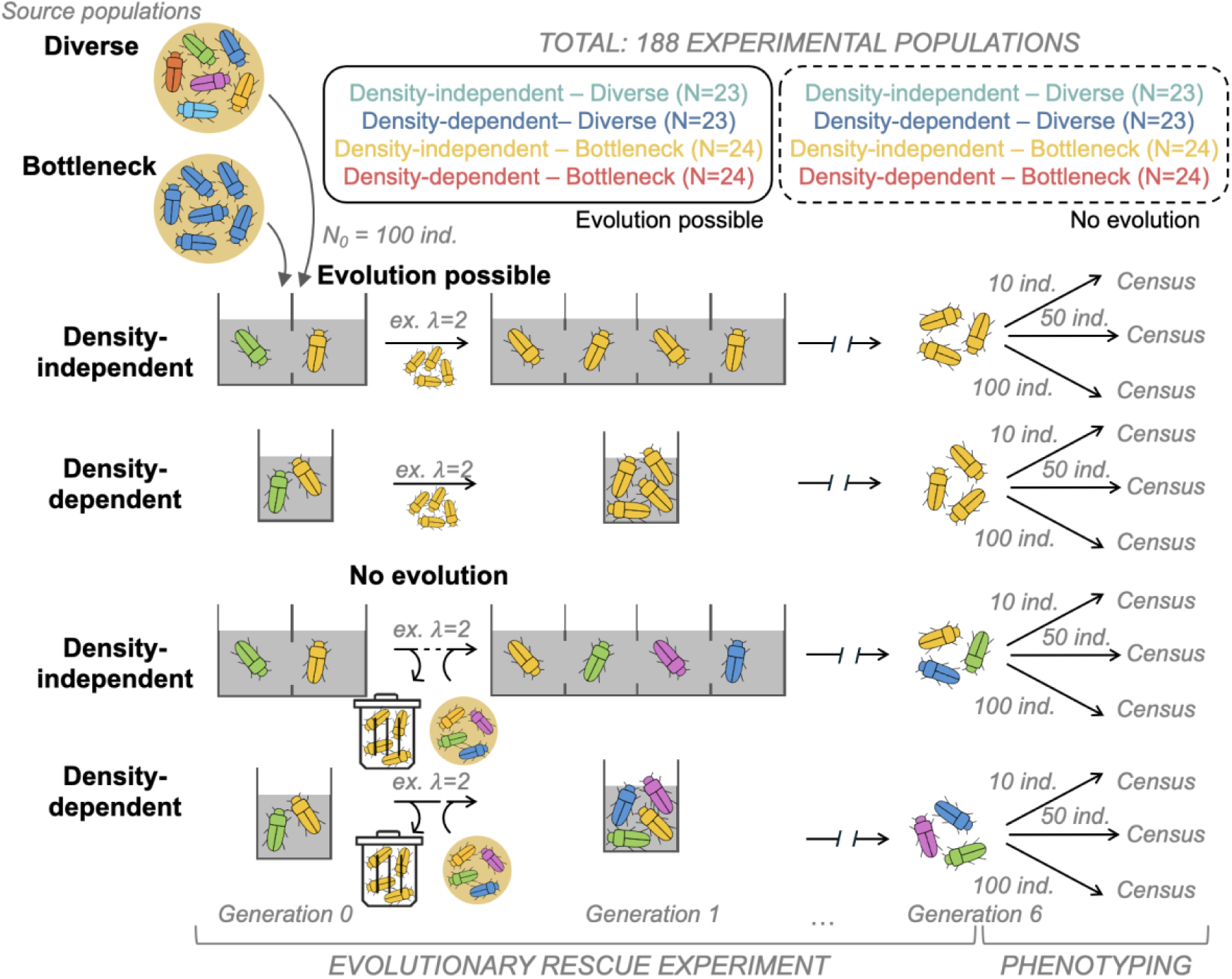
Experimental Design. The design of the factorial experiment manipulating, density dependence (density independent vs density dependent), genetic diversity (diverse vs. bottleneck), evolution (evolution possible vs. no evolution) giving eight treatment combinations (2×2×2). The 188 experimental populations were initiated with 47 distinct source populations (23 diverse and 24 bottlenecked populations) of T. castaneum, over four temporal blocks.

The 100 founders of each experimental population were placed into a challenging environment that was maintained consistently for six generations. The challenging environment was created by adding a pyrethroid insecticide, deltamethrin (DeltaDust, Bayer), to the standard medium at a concentration of 11.30 ppm (92.9% wheat flour, 4.9% brewer’s yeast and 2.2% DeltaDust). This environment produced average realized fitness in the first generation of exposure (at a test density of 50 individuals per patch) of 0.34 for diverse and 0.13 for bottlenecked populations (for comparison the average realized fitness on standard medium at a density of 50 throughout the experiment was 2.94 for diverse and 2.77 for bottlenecked populations). This environment was intended to impose a high risk of extinction across all treatments unless they were able to adapt to the stressful medium. We anticipated that adaptation would be likely, as resistance to deltamethrin mediated by cytochrome P450s is well-known in *T. castaneum* (Stuart *et al*. 1998, Zhu *et al*. 2010), and resistance can sweep through populations quickly (e.g., 12 bouts of selection can increase resistance nearly 100-fold; Khalid *et al*. 2023). Inheritance patterns in resistance suggest more than one gene is involved (Collins *et al*. 1998). Our approach allows for shifts in microbial communities or transgenerational or epigenetic changes to occur in response to selection in populations with a continuous line of descent (populations with evolution possible). Thus, these mechanisms may also contribute to observed phenotypic shifts. The strongly selective environment created by using deltamethrin allowed us to test the potential for evolutionary rescue under different ecological and genetic scenarios.

Each generation, we censused adults completely, then placed in patches with fresh challenging media for the 5-week life cycle described above. A population was considered extinct when the population size was 0 or 1 at the census.

To test the influence of density dependence on extinction and fitness, we compared experimental populations with low and constant density intended to approximate density independence with populations with naturally varying density. Populations in the density-independent treatment were maintained at 10 individuals/patch; as populations grew or declined, we increased or decreased the number of patches accordingly. Populations in the density-dependent treatment spent the whole experiment in a two-patch connected environment, regardless of population size. In the density-independent treatment because each patch contained 20 g of medium, resource availability was high and constant. Nonetheless, even at this low density of 10 individuals/patch some competition and cannibalism could potentially still occur. However, this low density would substantially reduce negative density dependence compared to the density-dependent treatment while avoiding positive density dependence such as Allee effects (e.g., difficulty in finding a sexual partner) that might have occurred if the density had been lower. We use the term “density-independent” for this low and constant density treatment, acknowledging that it approximates true density independence. It clearly contrasted with the density-dependent treatment, where density varied substantially over time, and drove strong competitive interactions and other forms of negative density dependence.

Where evolution was possible, the size of the populations under density independence increased rapidly during the experiment. For feasibility and logistical reasons, it was necessary to cap population size at 500 after complete census for the last two generations. Even though the population size was constrained, the density treatment was maintained for the whole experiment. Because density remained constant, we were able to project future population sizes based on observed growth rates. These projected values are conservative estimates, as the fitness of unrestricted (non-truncated) populations could potentially have increased even more rapidly due to their larger effective population sizes.

To test and control the influence of evolution, experimental populations were allowed to evolve (evolution-possible populations) or not (no-evolution control populations). In populations where evolution was possible there was a continuous line of descent, as is typical in experimental evolution. Individuals emerging from the challenging environment each generation comprised the next generation, allowing descent with modification (Darwin 1859). The dynamics of evolution-possible populations were due to demographic and adaptive effects. In no-evolution populations, the individuals emerging from the challenging environment each generation were replaced by the same number of individuals of the same age from the source population used to initiate the experimental population (as in Szucs *et al*. 2017). The source populations were maintained in parallel in the laboratory on standard medium with a population size of 600 individuals (density of 50 individuals per patch). This treatment prevented no-evolution populations from adapting to the novel environment. The dynamics of the no-evolution populations were therefore due to demographic effects but not selection. Comparison of no-evolution control populations with evolution-possible populations allows us to separate the adaptive effects (observed only in evolution-possible treatments) from purely demographic fluctuations in population size (present in both evolution-possible and no-evolution treatments).

### Final fitness assay

After five generations under these experimental conditions, the realized fitness of the experimental populations (evolution-possible and no-evolution) that did not go extinct was measured in a reciprocal transplant experiment (Kawecki & Ebert 2004, Kawecki *et al*. 2012). Specifically, directly after laying eggs to initiate the 6th generation, the 5th generation adults (now 36 days old) were used to measure realized fitness in the challenging environment at three densities (10, 50, and 100 individuals per patch). Because these beetles came directly from the experimental environment, this experiment does not control for potential epigenetic and microbial effects. Realized fitness was measured as the number of offspring produced per individual. The low density (10/patch) matches that used in the density-independent treatment. The intermediate density (50/patch) leads to negative density dependence and was the density used to initiate the density-dependent treatment in the evolutionary rescue experiment. The high density (100/patch) was commonly found in the density-dependent experimental populations.

For evolution-possible populations, the number of replicates depended on size of the experimental population. The no-evolution populations were quite small, and thus to have sufficient replication, we combined them with individuals from the corresponding lab source populations, which we maintained for 24 hours in the challenging environment at a density matching the evolutionary rescue experiment (35 days on standard medium initiated at 50 individuals/patch then 24 hours in the challenging medium at 10 individuals/patch for density-independent populations and 210 individuals/patches for density-dependent populations) to have enough individuals. *A posteriori*, we confirmed that the individuals coming directly from the experimental no-evolution populations (those that laid the 6th generation) or from the source populations (those that waited 1 day in the same experimental conditions) had the same pattern of realized fitness across the three densities (results not shown).

In total, because of differences in extinction, we had 6, 8, 11, and 20 surviving evolution-possible populations in our four density × diversity treatments. We had more surviving no-evolution populations and were also able to include extinct no-evolution populations using individuals from their source populations for a total of 138 populations. We quantified realized fitness of these populations at each of the three test densities (10, 50, 100). We additionally replicated within population using an average of 6.4 replicates per population at a density of 10, 4.5 replicates at a density of 50 and 1.4 replicates at a density of 100 (Table S1).

### Statistical analysis

All statistical models were fitted in R version 3.4.2 (R Core Team 2014). For all models, we estimated the significance of interactions and main effects by comparing a full model (i.e., with model terms of the same order) to a reduced model without the interaction or main effect of interest using a Likelihood Ratio Test (LRT). We used the full model describing the complete experimental design to estimate the magnitude and uncertainty of experimental effects and mean responses figures. To test specific hypotheses, we performed backward model selection by sequentially removing non-significant interactions using LRTs, while retaining all lower-order terms involved in significant interactions.

To understand how the risk of extinction in the challenging environment was influenced by density dependence, bottlenecks, and evolution, we used survival analysis. The response variable was the survival status (“surviving” or “extinct”) of each of the 188 populations across all six generations of the experiment. We estimated survival curves using the Kaplan–Meier method and compared survival dynamics across the 8 treatments using the log-rank test, implemented in the package *survival*. To evaluate the effects of explanatory variables on extinction risk, we used a Cox proportional hazards model, which estimates relative hazard rates, that is, the relative risk of extinction at any given time. In the proportional hazards model we used all variables and interactions of the factorial design of the experiment as fixed effects. Source population identity and temporal block were included as random effects. This mixed model was implemented with the package *coxme* (Therneau 2024). For visual representation, we expressed the probability of extinction during the experiment as 1 minus the Kaplan–Meier survival estimate.

To understand how the dynamics of the surviving populations were influenced by density dependence, bottleneck status, and evolution, we modeled population size through time using a Generalized Additive Mixed Model (GAMM ; Wood 2017) with a negative binomial distribution to account for overdispersion using the package *mgcv* (Wood 2022). Model diagnostics indicated substantial overdispersion when using a Poisson distribution (dispersion parameter = 11.01), which was effectively corrected by the negative binomial distribution (dispersion parameter = 1.06). The response variable was population size for all six generations of the experiment of each of the 91 populations that survived to the end of the experiment. We focused on surviving populations as our goal was to examine temporal dynamics specifically among populations that persisted long enough to potentially exhibit evolutionary rescue. We used projected population size of density-independent populations (see above) when population size was experimentally truncated. To allow for nonlinearity, we used a thin plate regression spline (Wood 2017). We used all variables and interactions of the factorial design of the experiment as well as generation (a continuous variable from 1-6) as fixed effects and source population identity and temporal block as random effects. This allowed the shape of the curve to vary by treatment over time. To test specific hypotheses, reduced models were compared to this full model using likelihood ratio tests with the *compareML* function of the package *itsadug* (Jacolien *et al*. 2022).

To determine how the intrinsic fitness of the surviving populations changed over time, including identifying when increases in fitness occurred, we fitted separate linear mixed-effects models for each of the four types of populations where evolution was possible in a 2×2 factorial design (diverse vs. bottlenecked evolution-possible populations under density-dependent or density-independent conditions). The response variable was the realized growth rate, ln(*N_t_*_+1_/*N_t_*), where *N_t_* was the population size at generation *t*, including all six generations of the 45 surviving evolution-possible populations. To estimate a breakpoint where the intrinsic fitness plateaued, realized growth rate was modeled as a piecewise linear function of generation. The breakpoint was estimated by profiling the likelihood function.

Population size (equivalent to density per box) in the previous generation was included as an explanatory variable for density-dependent treatments. The source population was modeled as a random effect. Confidence intervals were obtained via parametric bootstrapping. This approach corresponds to fitting a Ricker model for the density-dependent treatments, or an exponential model for the density-independent treatments. The Ricker model,

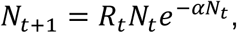

provides an accurate description of the biological processes occurring in this *Tribolium castaneum* model system (Melbourne & Hastings 2008), where *α* is the density-dependence parameter and *R_t_* is the intrinsic fitness at generation *t*. The linearized version,

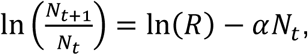

allows the model to be fitted by linear regression and the intrinsic growth rate, *r_t_* = ln(*R_t_*), to be estimated. Similarly, the linearized model for exponential growth is

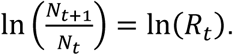

In addition, to determine how the final population size was influenced by density dependence, bottleneck, and evolution, we fitted a generalized linear mixed model, using the package *lme4* (Bates *et al*. 2015). The response variable was the population size (including integer values of projected population sizes) of each of the 91 populations that persisted to generation 6 (45 evolution-possible, 46 no-evolution). We used all variables and interactions of the factorial design of the experiment as fixed effects and source population identity and temporal block as random effects. Population size was modeled as a Poisson distribution with an observation-level random effect to account for overdispersion.

To further understand how the surviving populations had adapted to the challenging environment by the end of the study and how this depended on the density in which they were measured, we studied the realized fitness of each surviving population at different densities with a linear mixed model using the package *lme4* (Bates *et al*. 2015). The response variable was the realized fitness (i.e., number of offspring divided by the number of parents) measured experimentally at the three densities in the fitness assay. Realized fitness was modeled with a normal distribution (which modeled the residual variation better than log transformation or square root transformation). The model included density dependence, bottleneck, evolution, and their interactions as fixed effects, and source population identity and block as random effects. Additionally, we studied how these factors influenced the slope of the realized fitness as a function of density (continuous variable ranging from 0 to 100 but measured as three discrete values: 10, 50 and 100 individuals). By adding interactions between the discrete variables (density dependence, bottleneck, evolution) and the continuous variable (test density), we tested whether the slopes were significantly different between treatments.

## Results

### Extinction increased in density-dependent and bottlenecked populations

During the evolutionary rescue experiment, 97 of the 188 experimental populations maintained for 6 generations in a challenging environment went extinct. Extinctions were observed in all eight treatment combinations, ranging from a low of 4% in density-independent, diverse, no-evolution populations to a high of 88% in density-dependent, bottlenecked, no-evolution populations. Extinctions were observed in each generation of the experiment (Fig. S1).

Density dependence and a bottleneck both increased the risk of extinction. Populations able to evolve in density dependence had an extinction risk 18.1 times higher than populations evolving in density independence (CI 95% of the hazard ratio: [6.07, 53.9], *P-value* < 0.001). Bottlenecked populations had an extinction risk 15.4 times higher than diverse populations (CI 95% of the hazard ratio: [4.76, 49.7], *P-value* < 0.001). These effects were not additive and were reduced slightly when considered together: the risk of extinction was reduced by 0.18 for bottlenecked populations evolving in density dependence (CI 95% of the hazard ratio: [0.05, 0.58], *P-value* = 0.005) compared to adding the effects individually. This decrease is likely because bottlenecked populations remained small, so the effects of density dependence were weaker. We did not detect an effect of evolution on the risk of extinction as we had predicted or of its interactions with density dependence or bottleneck status (LRT *P-values* > 0.05). These findings were confirmed by the analysis of the probability of extinction, estimated as the probability that a population will not survive until the end of the experiment (Fig. 2). For instance, density dependence more than doubled the probability of extinction for diverse populations (Fig. 2a) but had a smaller effect on bottlenecked populations (Fig. 2b).

**Figure 2.**
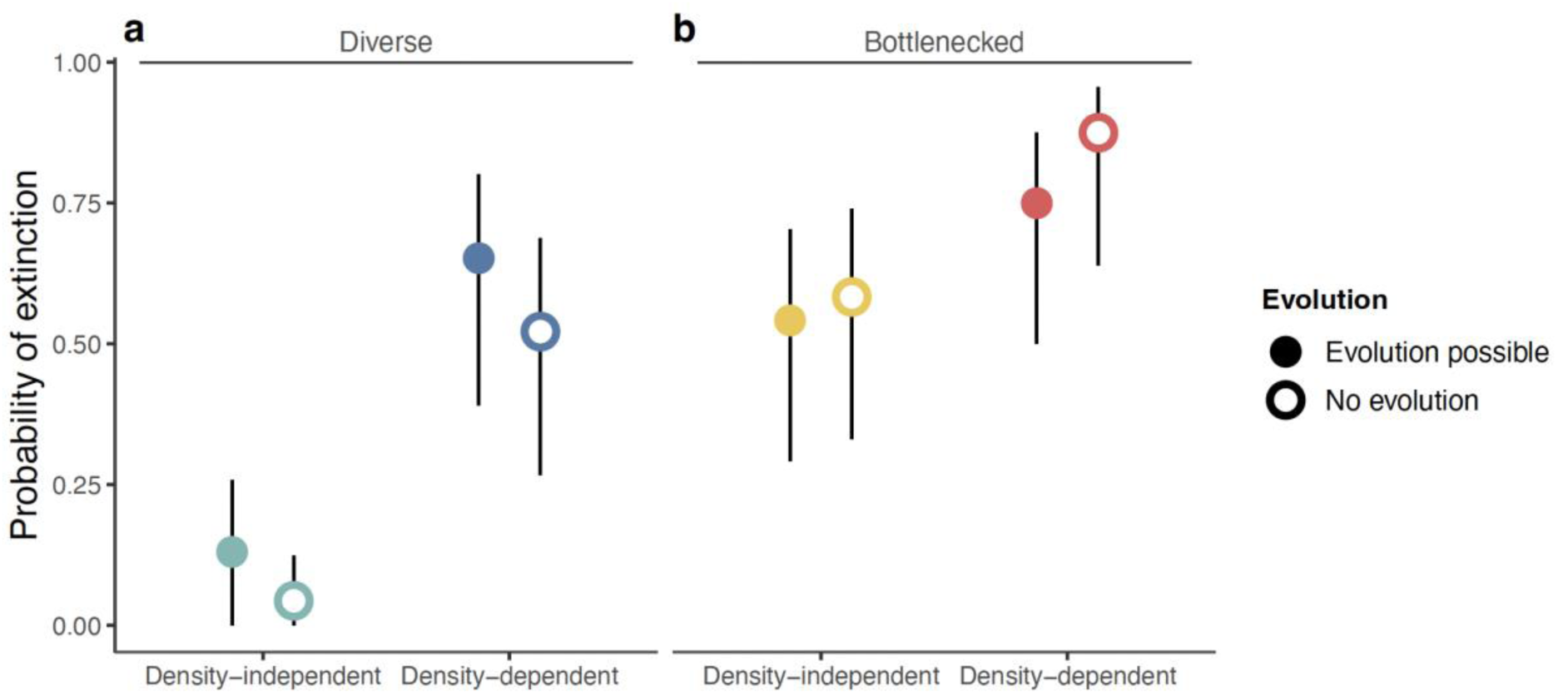
Probabilities of extinction showing that extinction was higher in bottlenecked and density-dependent populations. Error bars are 95% confidence intervals for the probabilities.

### Populations grew larger under density independence

We provide two visualizations of population dynamics: raw abundance data and mean abundances estimated from a generalized additive mixed model.

The raw data, presented on two scales, illustrate clearly that density dependence, bottleneck status, and evolution each shape population size through time (Fig. 3). Density independence enabled population growth, particularly in diverse, populations that could evolve (Fig. 3a-i, 3a-ii). In contrast, their no-evolution counterparts (Fig. 3e-i, 3e-ii) grew much less (Fig. 3a-i, 3a-ii), showing that response to selection was crucial for the dramatic population growth in density-independent, diverse populations. At generation 6, diverse, density-independent, populations that could evolve reached a mean size of 5,563 individuals while their no-evolution counterparts had a mean size of only 283 individuals (using data projected from growth rates where necessary, as described in the methods). The population size data also show that a previous bottleneck combined with density dependence restricted population growth substantially (Fig 3, looking by row, compare the last column with the previous three columns).

**Figure 3.**
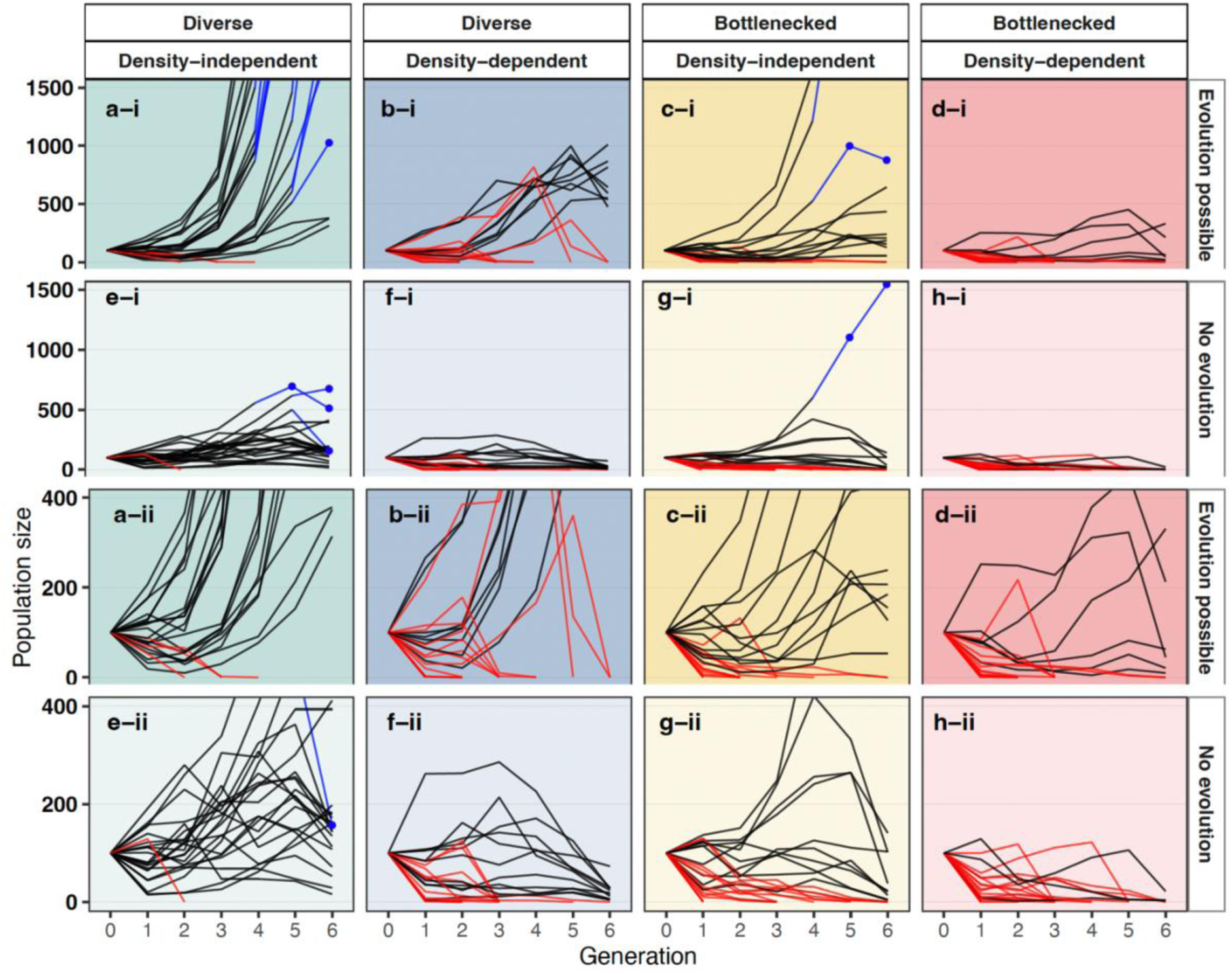
Size of experimental populations through time, showing that a population bottleneck and density dependence constrained population size and increased extinction risk, while density independence and evolution facilitated population growth. For easier comparison, the first two rows are on a scale of up to 1,500 individuals (a-i:h-i), while the last two rows repeat the same data but on a scale of up to 400 individuals (a-ii:h-ii). Red lines indicate populations that have gone extinct. Some density-independent populations grew too large to maintain in the incubators; in these cases, 500 individuals were used to produce the next generation, and their population size (shown in blue) was projected from their growth rate at constant low density. See methods for details.

We used a generalized additive mixed model (GAMM) to estimate the mean dynamics of surviving populations through time. Density dependence, bottlenecks and the evolution treatment each shaped these dynamics (LRTtriple interaction = 10.36, *P-value* < 0.001; Fig. 4). Diverse, populations that could evolve grew the most quickly, but density dependence limited the final size so that it was comparable to bottlenecked, density-independent populations. No-evolution populations were substantially smaller than their evolution-possible counterparts with density independence increasing final population size relative to density dependence (Fig. 4b, Fig. S3).

**Figure 4.**
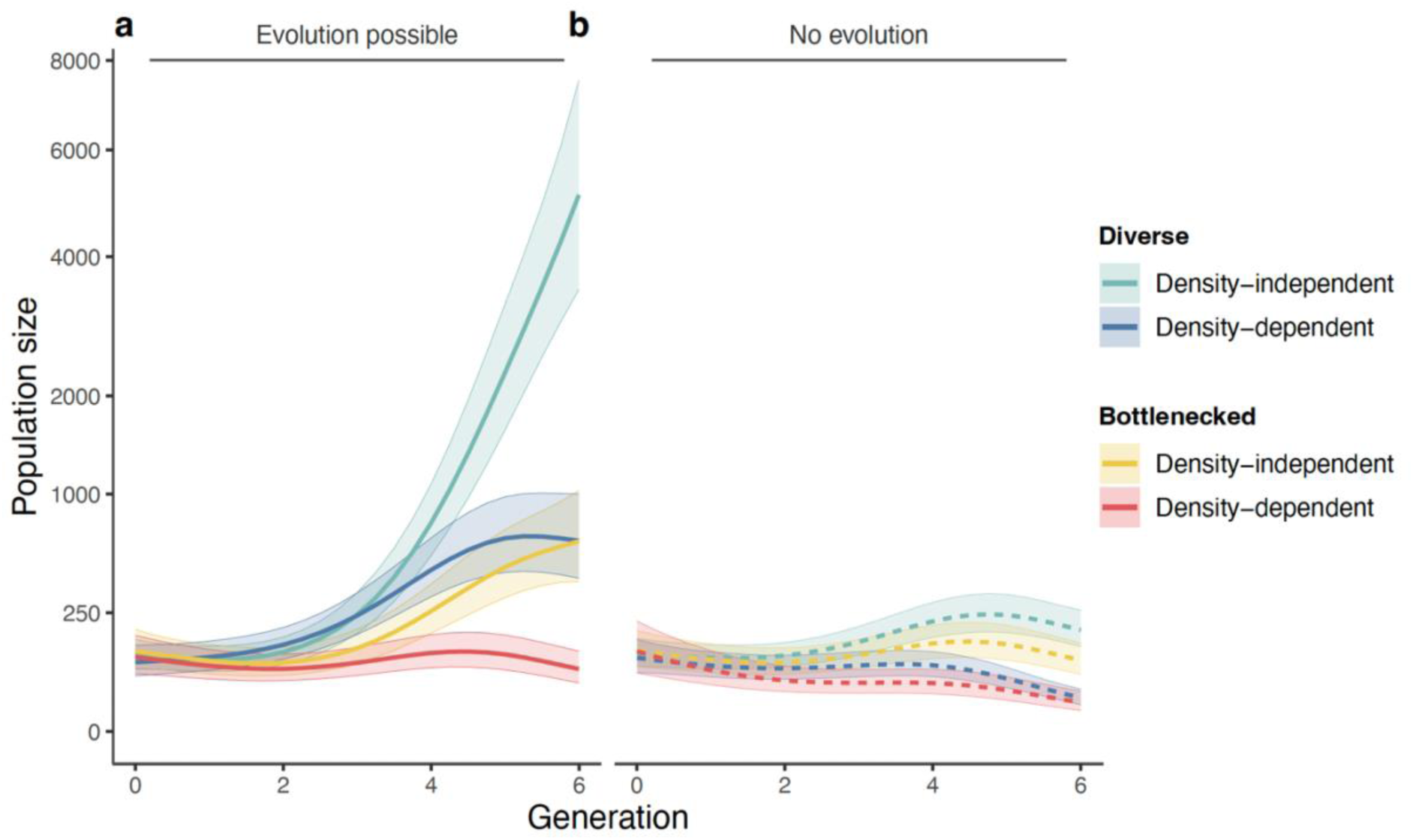
Trajectories of surviving populations through time, estimated by the generalized additive mixed model, showing that population sizes increased rapidly when diverse populations could adapt under density independence (panel a), and remained small when evolution was prevented, particularly under density dependence (panel b). The y-axis is on a square root scale to increase the visibility of small population sizes. The lines are estimates of the mean population size for each treatment from the model, averaged over the four temporal blocks and the 39 source populations (included in the model as random effects). Shaded areas are 95% confidence intervals.

### Intrinsic fitness increased for populations that could evolve, showing evidence consistent with adaptive evolution

The intrinsic growth rate (the natural log of intrinsic fitness) increased rapidly in populations where evolution was possible, with adaptation taking between 2.47 to 3.03 generations (Table S1). The rate of increase in growth rate (the slope of the line) was greater for diverse populations (Fig. 5, Table S2). In those diverse populations, density dependence led to a higher maximum intrinsic growth rate (rmax = 1.53, Fig. 5a) than density independence (rmax = 0.95, Fig. 5a). Bottlenecked populations achieved much lower intrinsic growth rates (rmax = 0.19 under density dependence and 0.46 under density independence).

**Figure 5.**
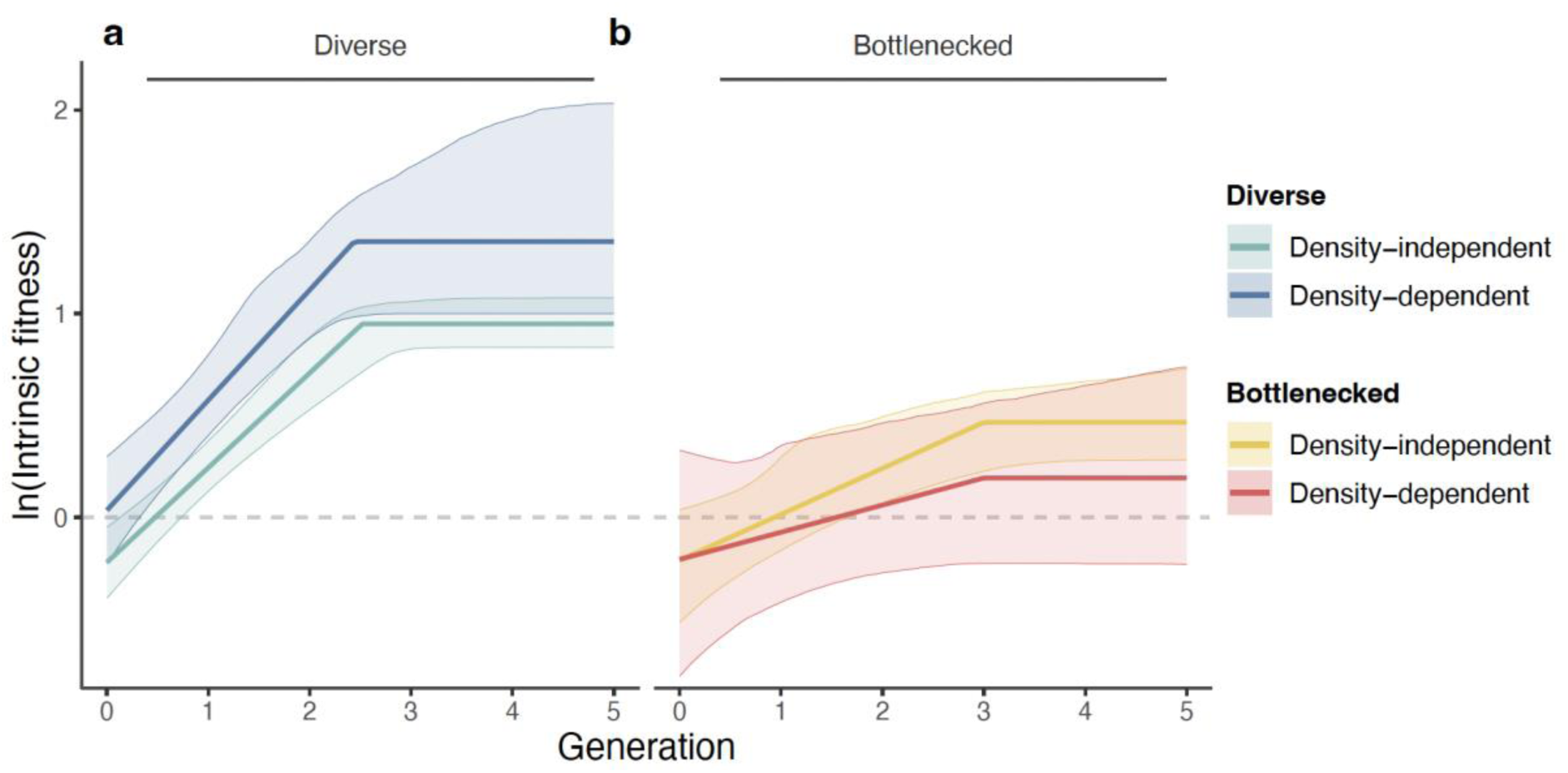
Evolution of intrinsic fitness through time for surviving populations, showing that populations adapted mainly during the first few generations and that the rate of adaptation was greater for diverse populations than bottlenecked ones. Among diverse populations, density dependence led to a higher intrinsic growth rate than density independence. The lines are estimates of the **changing** mean intrinsic growth rate (from piecewise linear models fitted to the data for each treatment separately). Shaded areas are 95% confidence intervals (parametric bootstrap). Ln(intrinsic fitness) above the grey dashed line indicates where populations could grow from low density.

### Realized fitness at the end of the experiment confirmed increased fitness in populations allowed to evolve

Realized fitness of the surviving populations at controlled densities confirm that populations where evolution was possible showed evidence consistent with adaptative evolution, having higher realized fitness in the challenging environment than the no-evolution populations (Fig. 6, Table S3). Realized fitness increased most in diverse populations (Fig. 6). Realized fitness was reduced at higher densities, particularly for bottlenecked populations, showing their greater sensitivity to negative density dependence (LRTdensity x evolution x bottleneck interaction: χ² = 6.36, *P-value* = 0.01). Populations that evolved under density dependence and density independence showed similar patterns (Fig. 6, LRTintercept: χ² = 0.12, *P-value* = 0.73, LRTslope: χ² = 0.39, *P-value* = 0.53 and always P-values > 0.33 when considered in interaction with other variables). Nevertheless, at high density, realized fitness of density-dependent populations appeared slightly higher than density-independent populations (Fig. 6). This pattern is consistent with adaptation to higher density, more efficient selection, or both in density-dependent populations, although further evidence would be needed to confirm this.

**Figure 6.**
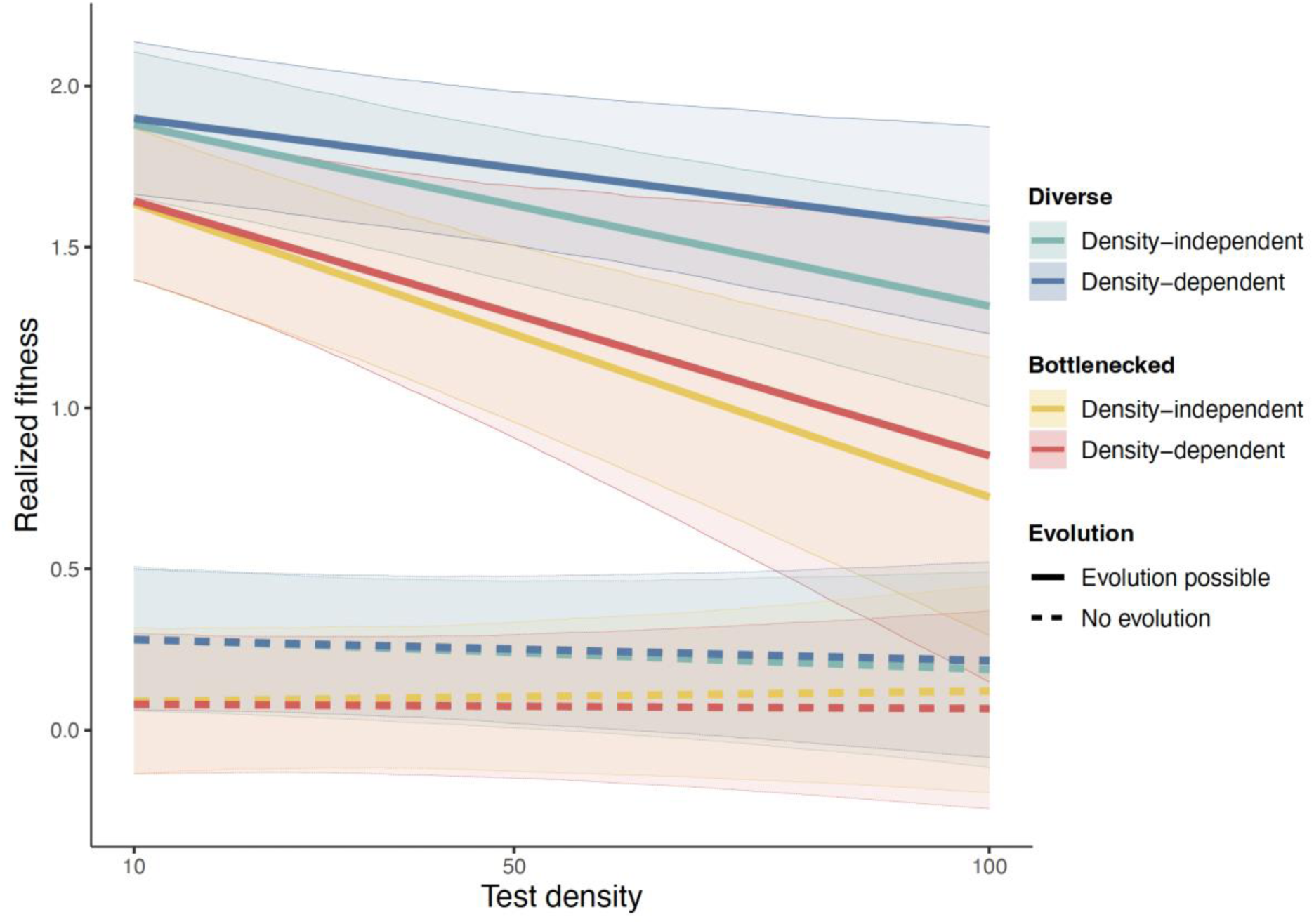
Mean realized fitness of surviving populations after the rescue experiment, as a function of test density. The lines are estimates from the linear mixed model of the mean realized fitness for each treatment, averaged over the four temporal blocks and the 47 source populations (included in the model as random effects). The model included all aspects of the experimental design, including all interactions of the treatment factors. Shaded areas are 95% confidence intervals (parametric bootstrap).

## Discussion

Here, we show experimentally that density dependence greatly increases the extinction risk of populations in challenging environments but nonetheless does not prevent the evolution of increased fitness in populations that are able to persist.

Additionally, the effects of density dependence are influenced by the genetic diversity of the population. Populations that have experienced bottlenecks in size, which can both reduce genetic diversity and increase genetic load, have a yet higher risk of extinction under density dependence, and a much lower ability to adapt to environmental change. Our results support theory (Nordstrom *et al*. 2023; Osmond & de Mazancourt 2013) as they are consistent with selection under density dependence leading to higher fitness among surviving populations, despite a higher extinction rate. Density independence enabled increased population sizes, a result that in itself is intuitive. The remarkable feature of this finding is that the increase in population size depended on evolution, showing the eco-evolutionary nature of population growth. Populations allowed to evolve under density independence were not able to increase exponentially or continuously, and populations with limited evolutionary potential (bottlenecked populations in the evolution-possible treatment) grew, but only slowly. Overall, our study highlights the crucial importance of both preserving genetic diversity and increasing habitat size and resource density to relieve density dependence for populations of conservation concern.

Density dependence increases extinction without necessarily limiting adaptation. In our experiment, density dependence had a strong impact on the trajectory of populations and their ability to persist over time. Density dependence increased the risk of extinction and suppressed growth of populations throughout the experiment. These findings disagree with theory that assumes populations under density dependence and density independence are adapted to those density conditions and thus start with equivalent realized fitness, which then leads to ecological release from negative density dependence when population size decreases (Reed *et al*. 2015; Vinton & Vasseur 2020) and instead align with theoretical frameworks that start with equivalent intrinsic fitness and thus density dependence reduces vital rates (Chevin & Lande 2010; Nordstrom *et al*. 2023).

Fundamentally, outcomes in nature may thus depend upon the degree to which, and the mechanisms by which, populations adapt to density dependence prior to detrimental environmental change. If density dependence generally reduces vital rates, natural populations adapting to drastic environmental change while experiencing negative density dependence may suffer, increasing their risk of extinction. Given that density-dependent dynamics are widespread in natural populations (Bonenfant *et al*. 2009), it is crucial to consider these effects in the management of populations that could be at risk of extinction when environments change.

The adaptive potential of populations that experienced a bottleneck was substantially reduced, particularly under density dependence. The combination of bottlenecks and density dependence was nearly as detrimental to the ability of populations to adapt and grow over time as being unable to evolve at all. Bottlenecks dramatically reduced the advantages of populations allowed to evolve under density independence. Where evolution was possible, surviving bottlenecked populations under density independence reached sizes similar to diverse populations under density dependence. Furthermore, even though the bottlenecked populations that persisted had increased fitness at the end of the experiment, they never reached the fitness of the diverse populations. This is because bottlenecked populations have a combination of both lower diversity and higher genetic load (Bertorelle *et al*. 2022a; Frankham *et al*. 1999), which makes them less able to adapt and therefore less able to take advantage of the density-independent conditions. These findings are of particular significance in the context of conservation, as populations facing extinction frequently experience both bottlenecks and restricted habitat sizes (Bouzat 2010).

For instance, Stoffel et al. examined 30 species of pinnipeds and demonstrated that the three species most significantly impacted by a bottleneck, resulting from hunting or habitat loss, had been designated as endangered (Stoffel *et al*. 2018). Our findings emphasize that bottlenecked populations are particularly vulnerable to extinction following detrimental environmental change when density-dependent effects are considered. In addition, density dependence and bottlenecks can be associated in nature, particularly when restricted habitat size leads to both a bottleneck and strong density dependence. As bottlenecks and density dependence are common in natural populations, our findings could help explain why adaptation to sudden environmental change is not commonly observed in nature (Pujol *et al*. 2018).

The clear differences in evolutionary trajectories between populations that could evolve and no-evolution populations are consistent with evolutionary adaptation to the novel environment, particularly given the known ability of *T. castaneum* to evolve resistance to deltamethrin (Zhu *et al*. 2010, Stuart *et al*. 1998). Furthermore, differences between genetically diverse and bottlenecked populations (which differ in standing genetic variation), strongly support adaptive evolution as a major driver of the patterns we observed. Additionally, maternal transgenerational effects and shifts in beetle microbiomes could contribute to the patterns we see. Indeed, all three of these mechanisms could be acting. Evidence from other systems suggests a potential role for the microbiome in particular. For example, deltamethrin has known toxic effects on microbial communities (Farghaly *et al*. 2013), and mosquitos differing in resistance to deltamethrin differ in some components of their gut microbiomes (Sun *et al*. 2024). Even more direct evidence that microbes can potentially play a role in resistance comes from bees, where sterilized bees (without gut microbiomes) perform better on exposure to deltamethrin if they are inoculated with certain strains of bacteria (Dong *et al*. 2022).

Further research could help determine which mechanisms explain our findings. For example, a transgenerational common garden experiment, and crossing of individuals from populations that could and could not evolve would help to determine the roles of transgenerational effects and adaptive evolution.

Additionally, genomic data on both the beetles and their microbiomes could help determine whether the observed phenotypic shifts correspond to changes in allele frequencies, microbial communities, or both. Such studies would be valuable in evaluating possible contributions of non-genetic factors in evolutionary rescue.

Our experimental design may have reduced extinction rates in no-evolution populations, leading to extinction probabilities similar to populations that were able to evolve. Individuals introduced each generation originated from the ancestral environment (without the toxin) and thus were not adapted to the stressful environment. However, the benign maternal environment may have enhanced reproduction and larval performance in the stressful environment, as in other studies (Olazcuaga *et al*. 2023; Van Allen & Rudolf 2013). This would potentially prevent populations from declining to sizes where they are at risk of extinction due to demographic stochasticity. However, without the ability to adapt, the no-evolution populations remain small, while populations that could evolve become considerably larger. Consequently, it is likely that if natural populations are unable to evolve at all, they may have a higher risk of extinction than demonstrated by the no-evolution populations in this experiment.

The long-term performance of populations after evolutionary rescue remains unclear. Our experiment demonstrates that if populations are able to persist and adapt to severe environmental change, density dependence may improve their fitness by increasing the strength of selection, particularly if they have adequate genetic diversity. Whether such a fitness boost lasts is unknown, as our experiment was limited to six generations. In the future, it would be interesting to re-examine these dynamics over an extended period for two key reasons. First, having passed through low population sizes, populations might have accumulated genetic load (Bertorelle *et al*. 2022b), which could reduce their persistence over the long term (Dussex *et al*. 2023). Second, the suppressing effects of high density increase in strength again as population size increases with adaptation. Consequently, because diverse populations are better able to adapt, they will experience stronger effects of density dependence in the long term. Few studies have examined the long-term persistence of populations following evolutionary rescue, which would be particularly relevant in the context of density dependence.

These findings have clear broader implications for managers of natural populations, highlighting the importance of both genetic diversity and habitat size. While even the bottlenecked populations were able to adapt to the stressful environment, the diverse populations attained much higher fitness. Thus, in natural populations known to or likely to have low diversity, genetic diversity should be enhanced by active management (Ralls *et al*. 2018, 2020). This should both reduce extinction risk and improve the ability of populations to adapt to changing environments. This recommendation is in alignment with the large literature on genetic rescue (the assisted movement of individuals among populations to enhance diversity and increase fitness), which can substantially beneficial populations (Frankham 2015; Ralls *et al*. 2020). We agree with others (Hedrick & Garcia-Dorado 2016; Whiteley *et al*. 2015) that masking genetic load and enhancing diversity is of paramount importance. However, our results show that care must be taken in assisted migration and genetic rescue to not increase population density too substantially, or risk increasing negative density dependence.

Along with genetic diversity, habitat size and quality should remain a focus in management and conservation efforts. Alleviating negative density dependence by enhancing either habitat size or resource availability should substantially aid populations, even bottlenecked or inbred ones. Evidence from natural populations shows that challenging environmental conditions are all the more detrimental to populations when they are close to their carrying capacity and density dependence is high (e.g., in feral sheep (Coulson *et al*. 2001) and Alpine ibex (Jacobson *et al*. 2004)). However, as challenging as it is to move organisms to enhance diversity, increasing habitat size and quality is likely to be more challenging. Nonetheless, our findings illustrate that reducing density dependence by increasing habitat size can be powerful in alleviating extinction risk, and we hope these finding can help support the political and social decisions needed to provide resources for habitat improvement.

To conclude, negative density dependence is common in natural plant and animal populations, often in the form of intraspecific competition (Adler *et al*. 2018; Bretagnolle *et al*. 2008; Grossman & Simon 2020). Here, we show experimentally for the first time that density dependence dramatically influences the ability of populations to persist in challenging environments, reducing the likelihood of evolutionary rescue.

Our results provide both challenges and hopes for the management of natural populations. The smaller size of populations experiencing density dependence clearly places them at risk of extinction, and alleviating density dependence through habitat improvements is challenging. On the more hopeful side, even populations in strongly density-dependent conditions can adapt to rapid environmental change, particularly if they harbor adequate genetic diversity. Adaptive evolution thus can counter some of the effects of density dependence, buying time for habitat restoration.

## Supporting information

SupMat

## Acknowledgements

We thank the excellent team of undergraduate students who participated with data collection: H. Degan, M. Hayes, L. Mendoza, D. Moore, R. McMillan, N. Namba, H. Nolan, P. Harper, F. Stevens, M. Wallace, B. Weigand and S. Wright; as well as Jorge Rivera-Gonzalez and Lily Durkee for any laboratory support. Funding for this research was provided by the US National Science Foundation (DEB-1930650 to R.A.H.; DEB-1930222 to B.A.M.).

## Notes

### Competing Interest Statement

The authors have declared no competing interest.

### Summary of Updates

We have added information in the methods and also a dedicated section in the Discussion that highlights the limitations of the study and explicitly outlines what additional approaches (e.g., molecular/genomic evidence) would be required to definitively confirm evolutionary change.

