## Supplementary material for "Density dependence during evolutionary rescue increases extinction risk but does not prevent adaptation": SupMat

### Density dependence impedes evolutionary rescue

#### Supplementary Information

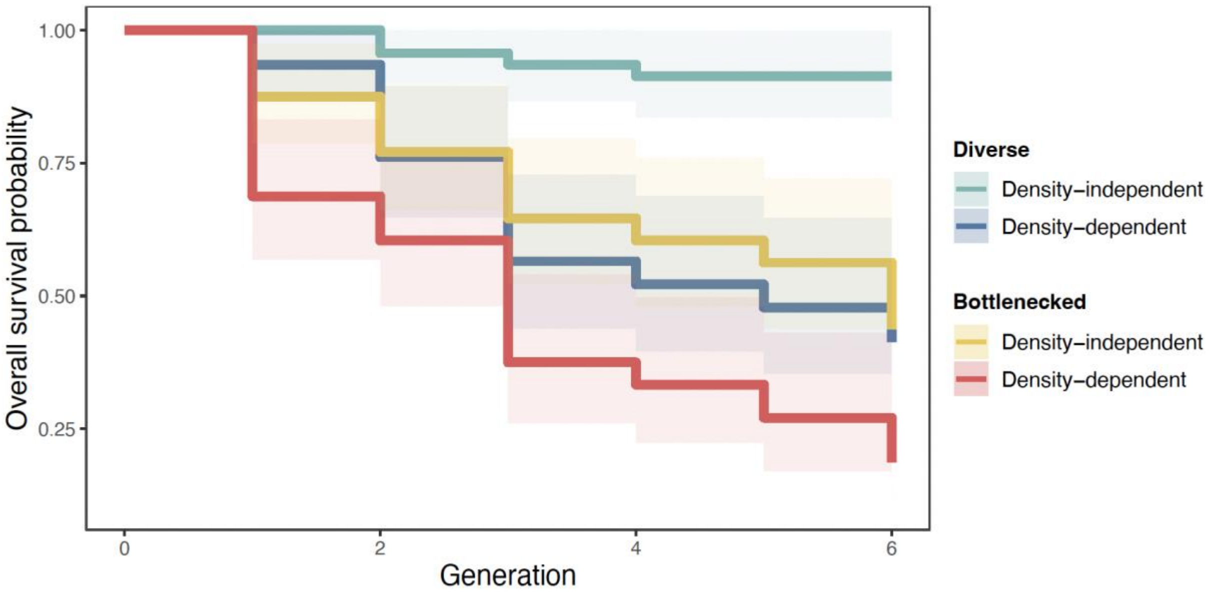


***Figure S1. Survival dynamics estimated by the best-fit survival model****. Survival probabilities were estimated by the best-fit survival model, where the Evolution-possible and No-evolution treatments are merged due to a lack of evidence of significant differences. Shaded areas are 95% confidence intervals for the probabilities.*

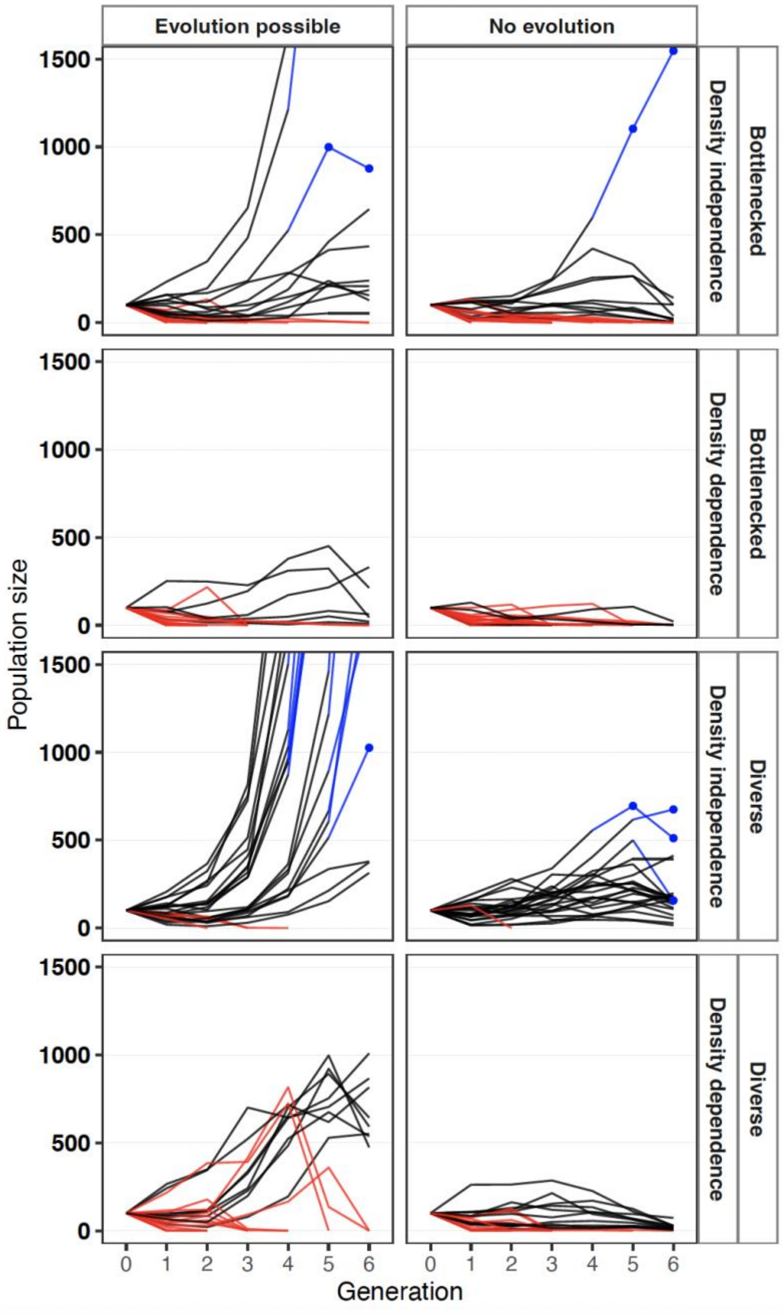


***Figure S2. Size of experimental populations through time, illustrating how selection decreases extinction risk, and how density independence and genetic diversity with evolution, facilitate population growth.*** *The scale of the y-axis is fixed at a maximum of 1500 individuals to facilitate comparison between Evolution-possible and No-evolution populations, and the lettering is consistent with Figure 1 (e.g., Fig. 2a and Fig. S2a contain the same data on different scales). Red lines indicate populations that went extinct. The blue dots and blue lines indicate projection of population size based on the growth rate of that generation, rather than the census population size.*

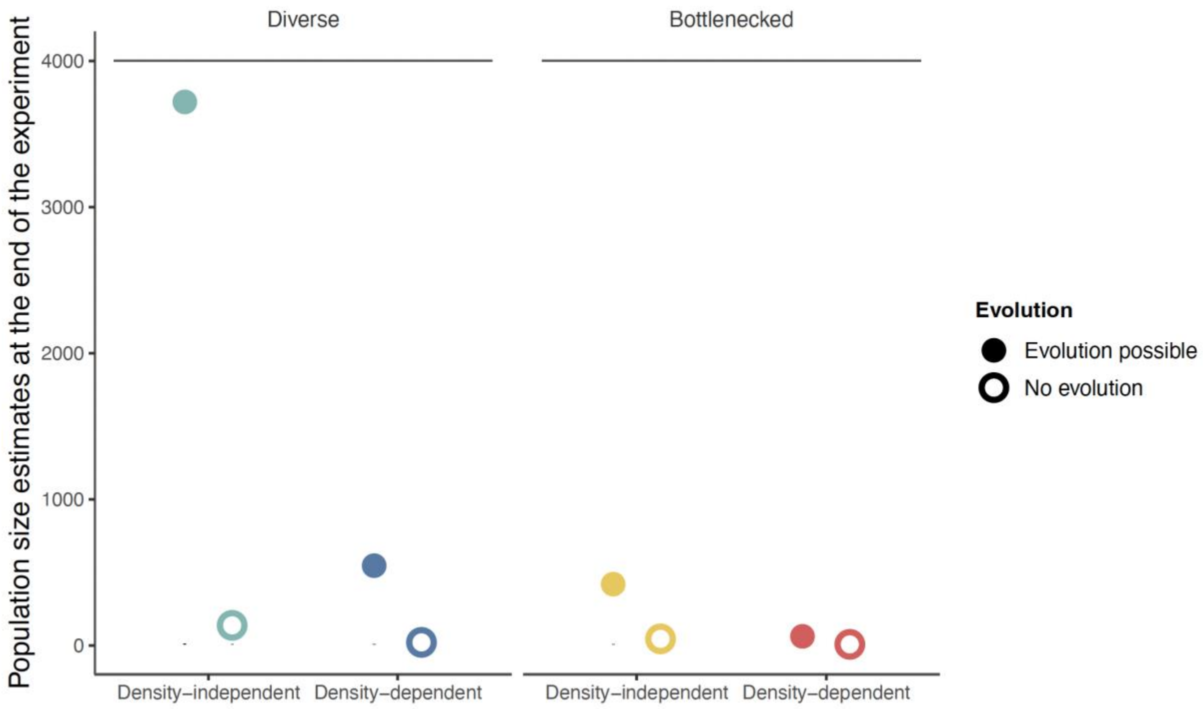


***Figure S3. Final population size of surviving populations estimated at the end of the experiment for each combination of treatments.*** *The final population size were estimated by the best-fit model. Error bars are 95% confidence intervals computed by bootstrapping to consider variation due to random effects (source population, temporal blocks, and overdispersion). These errors are not visible because of the scale of the y-axis. The data used for this analysis was the estimates of population size based on the growth rate of that generation, rather than the census population size, at the 6th generation of the experiment where the density of the parents was not experimentally controlled.*

The analysis of the final population size confirmed that no-evolution and bottlenecks both decreased the final size of the population, and their effects were not additive ($LRT_{evolution\ x\ bottleneck\ interaction}$: $\chi^2 = 5.46$, *P-value* = 0.019, Fig. S3).

***Table S1 Number of replicates for the fitness assay.*** *Treatments vary by density dependence, evolutionary history and selection treatments.*

| Treatment | 10ind | 50ind | 100ind |
|---|---|---|---|
| Diverse Density-independent Evolution-possible | 123 | 58 | 32 |
| Bottlenecked Density-independent Evolution-possible | 64 | 18 | 13 |
| Diverse Density-independent No-evolution | 145 | 75 | 30 |
| Bottlenecked Density-independent No-evolution | 147 | 84 | 24 |
| Diverse Density-dependent Evolution-possible | 74 | 29 | 34 |
| Bottlenecked Density-dependent Evolution-possible | 26 | 9 | 3 |
| Diverse Density-dependent No-evolution | 149 | 85 | 25 |
| Bottlenecked Density-dependent No-evolution | 146 | 94 | 25 |

***Table S2 Maximum estimated of the intrinsic growth rate (ln(intrinsic fitness))*** ***$r_{max}$ with associated 95% confidence intervals (CI) and corresponding generation for each experimental treatment.*** *Treatments vary by density dependence and evolutionary history.*

| Treatment | Time | $r_{max}$ | CI_lower | CI_upper |
|---|---|---|---|---|
| Diverse Density-independent | 2.53 | 0.95 | 0.72 | 1.03 |
| Bottlenecked Density-independent | 3.03 | 0.47 | 0.23 | 0.62 |
| Diverse Density-dependent | 2.47 | 1.35 | 0.99 | 1.57 |
| Bottlenecked Density-dependent | 3.03 | 0.19 | -0.23 | 0.56 |

***Table S3 Estimate of the realized fitness with associated 95% confidence intervals (CI) for each density tested and each experimental treatment.***

*Treatments vary by density dependence, evolutionary history and selection.*

| Treatment | Density | Realized fitness | CI-lower | CI-upper |
|---|---|---|---|---|
| Diverse Density-independent Evolution-possible | 10 | 1.88 | 1.65 | 2.11 |
| Diverse Density-independent Evolution-possible | 50 | 1.63 | 1.39 | 1.86 |
| Diverse Density-independent Evolution-possible | 100 | 1.32 | 1.01 | 1.63 |
| Bottlenecked Density-independent Evolution-possible | 10 | 1.63 | 1.39 | 1.87 |
| Bottlenecked Density-independent Evolution-possible | 50 | 1.23 | 0.95 | 1.51 |
| Bottlenecked Density-independent Evolution-possible | 100 | 0.72 | 0.29 | 1.16 |
| Diverse Density-independent No-evolution | 10 | 0.28 | 0.05 | 0.51 |
| Diverse Density-independent No-evolution | 50 | 0.24 | 0.01 | 0.47 |
| Diverse Density-independent No-evolution | 100 | 0.19 | -0.11 | 0.5 |
| Bottlenecked Density-independent No-evolution | 10 | 0.09 | -0.13 | 0.31 |
| Bottlenecked Density-independent No-evolution | 50 | 0.1 | -0.12 | 0.34 |
| Bottlenecked Density-independent No-evolution | 100 | 0.12 | -0.19 | 0.44 |
| Diverse Density-dependent Evolution-possible | 10 | 1.9 | 1.67 | 2.14 |
| Diverse Density-dependent Evolution-possible | 50 | 1.75 | 1.51 | 1.99 |
| Diverse Density-dependent Evolution-possible | 100 | 1.55 | 1.24 | 1.88 |
| Bottlenecked Density-dependent Evolution-possible | 10 | 1.64 | 1.38 | 1.89 |
| Bottlenecked Density-dependent Evolution-possible | 50 | 1.29 | 0.89 | 1.7 |
| Bottlenecked Density-dependent Evolution-possible | 100 | 0.85 | 0.14 | 1.58 |
| Diverse Density-dependent No-evolution | 10 | 0.28 | 0.06 | 0.5 |
| Diverse Density-dependent No-evolution | 50 | 0.25 | 0.02 | 0.48 |
| Diverse Density-dependent No-evolution | 100 | 0.21 | -0.09 | 0.53 |
| Bottlenecked Density-dependent No-evolution | 10 | 0.08 | -0.15 | 0.31 |
| Bottlenecked Density-dependent No-evolution | 50 | 0.07 | -0.16 | 0.3 |
| Bottlenecked Density-dependent No-evolution | 100 | 0.07 | -0.25 | 0.38 |
